## Supplementary Figures S1 to S9 for "Accurate Virus Identification with Interpretable Raman Signatures by Machine Learning"

**Corresponding Author(s):**

Sharon Xiaolei Huang

**This PDF file includes:**

Figures S1 to S9

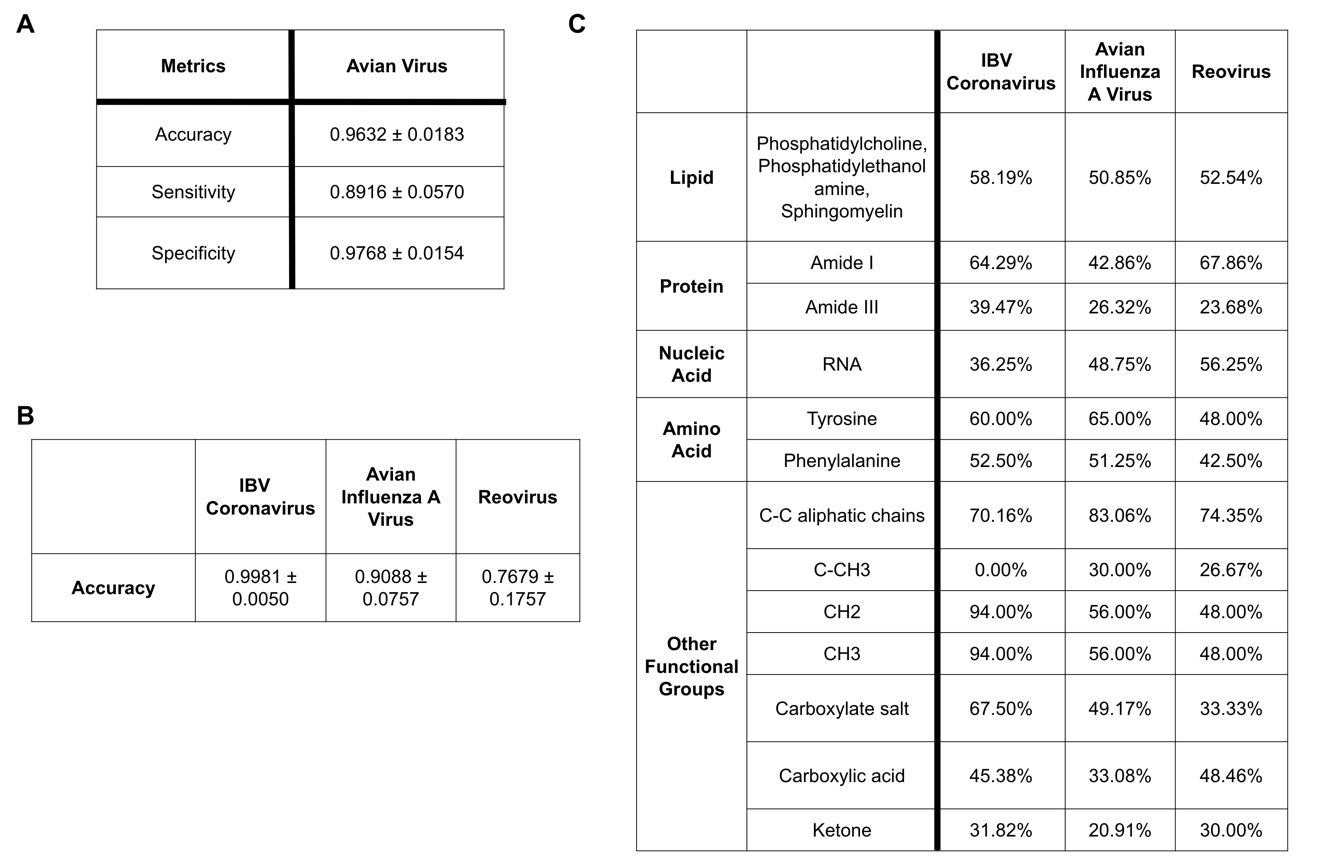

Fig. S1. A. The CNN classification performance of Avian viruses on three metrics (Accuracy, Sensitivity and Specificity); B. The CNN classification accuracy for each type of Avian virus; C. Matching scores between Raman ranges important for identifying Avian viruses using ML and Raman peak ranges of biomolecules.

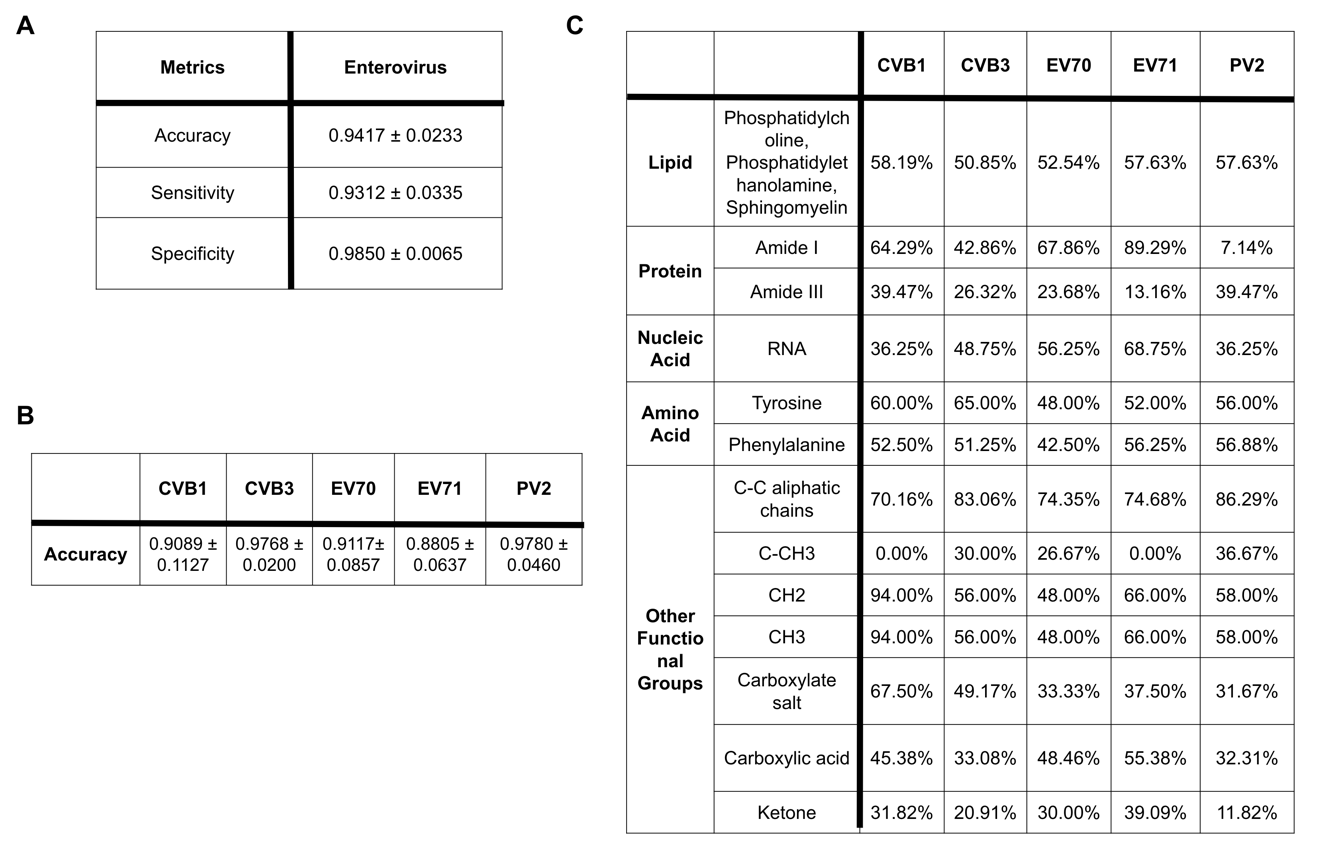

Fig. S2. A. The CNN classification performance of Enteroviruses on three metrics (Accuracy, Sensitivity and Specificity); B. The CNN classification accuracy of each type (subtype) of Enterovirus; C. Matching scores between Raman ranges important for identifying Enteroviruses using ML and Raman peak ranges of biomolecules.

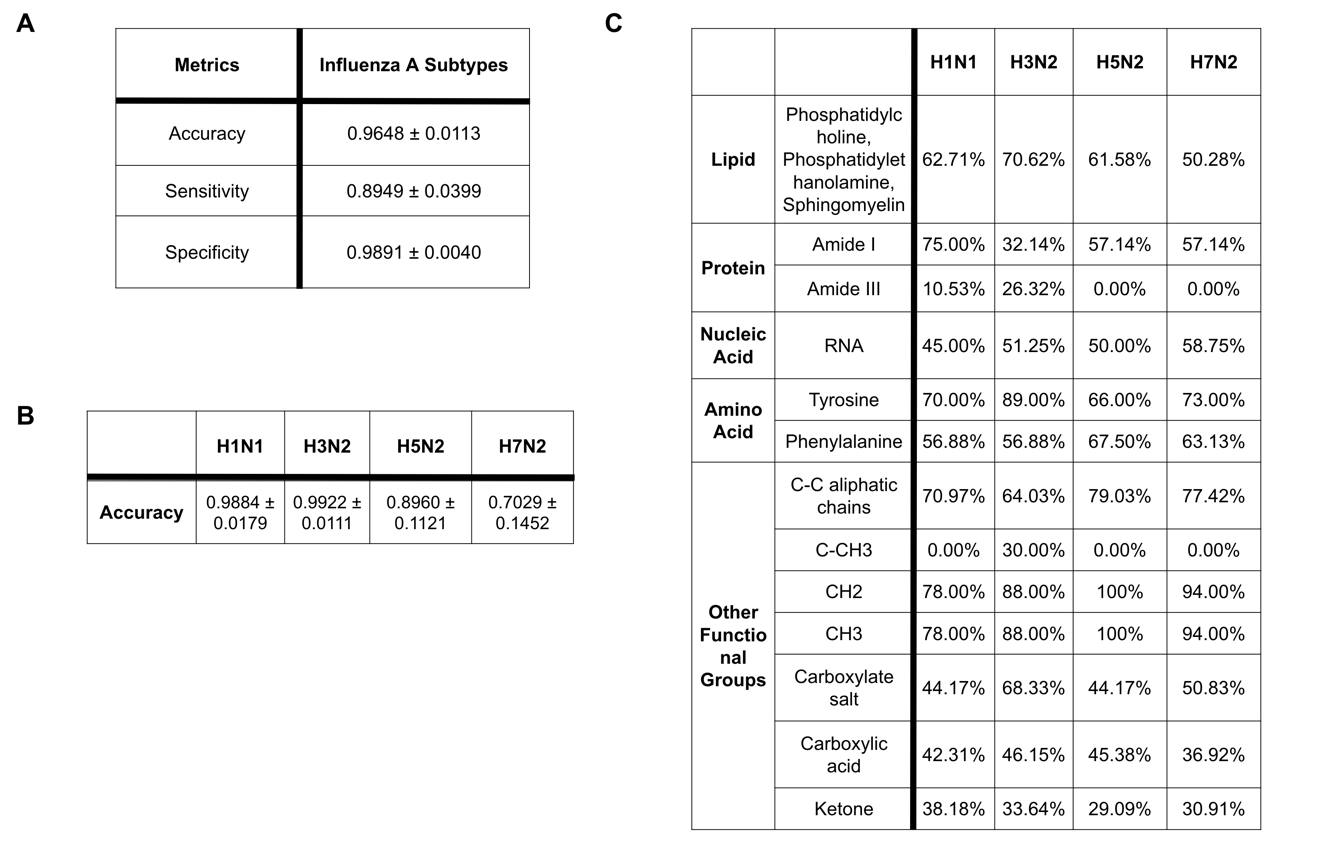

Fig. S3. A. The CNN classification performance of Influenza A virus subtypes on three metrics (Accuracy, Sensitivity and Specificity); B. The CNN classification accuracy of each subtype of Influenza A virus; C. Matching scores between Raman ranges important for identifying Influenza A subtypes using ML and Raman peak ranges of biomolecules.

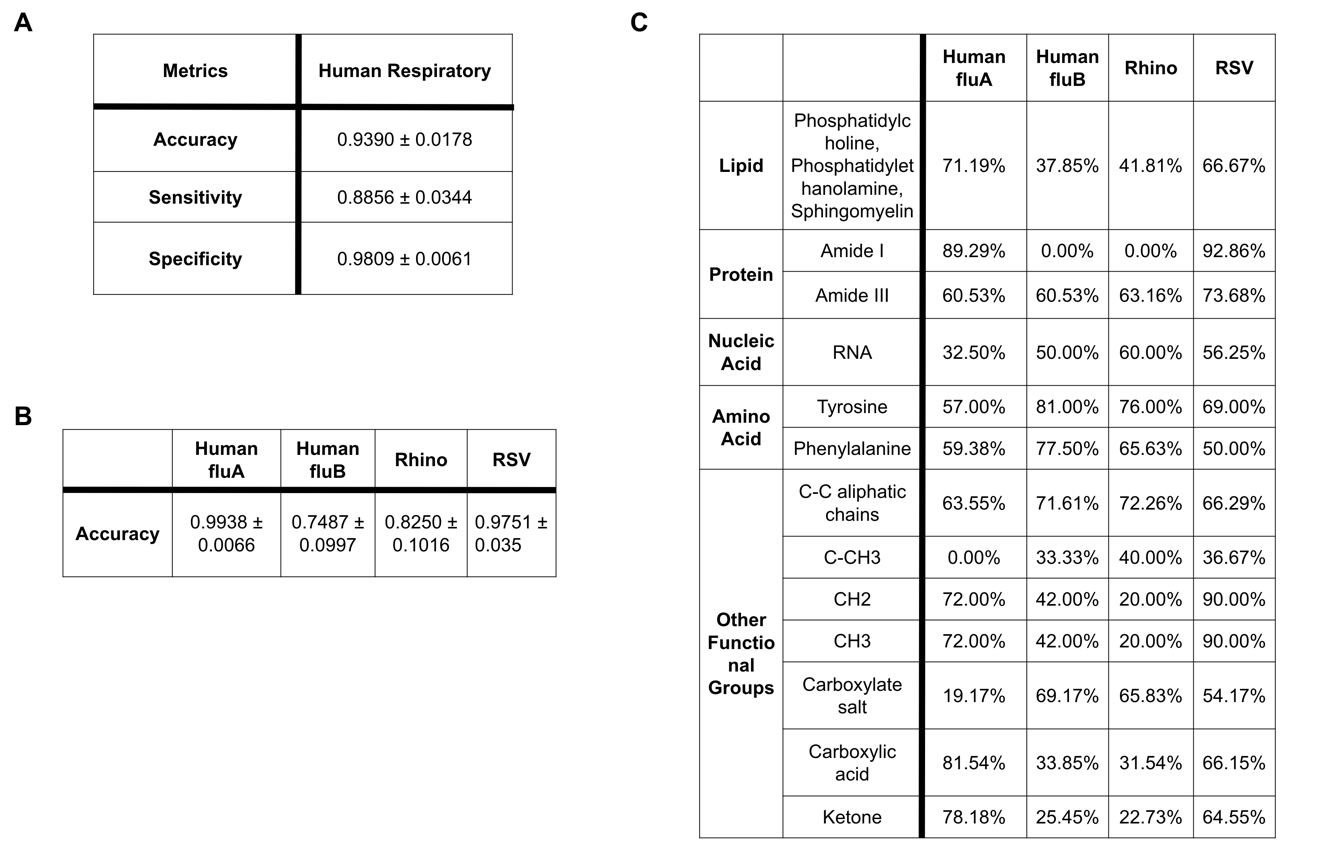

Fig. S4. A. The CNN classification performance of Human Respiratory viruses on three metrics (Accuracy, Sensitivity and Specificity); B. The CNN classification accuracy for each type of Human Respiratory virus; C. Matching scores between Raman ranges important for identifying different types of Human Respiratory viruses using ML and Raman peak ranges of biomolecules.

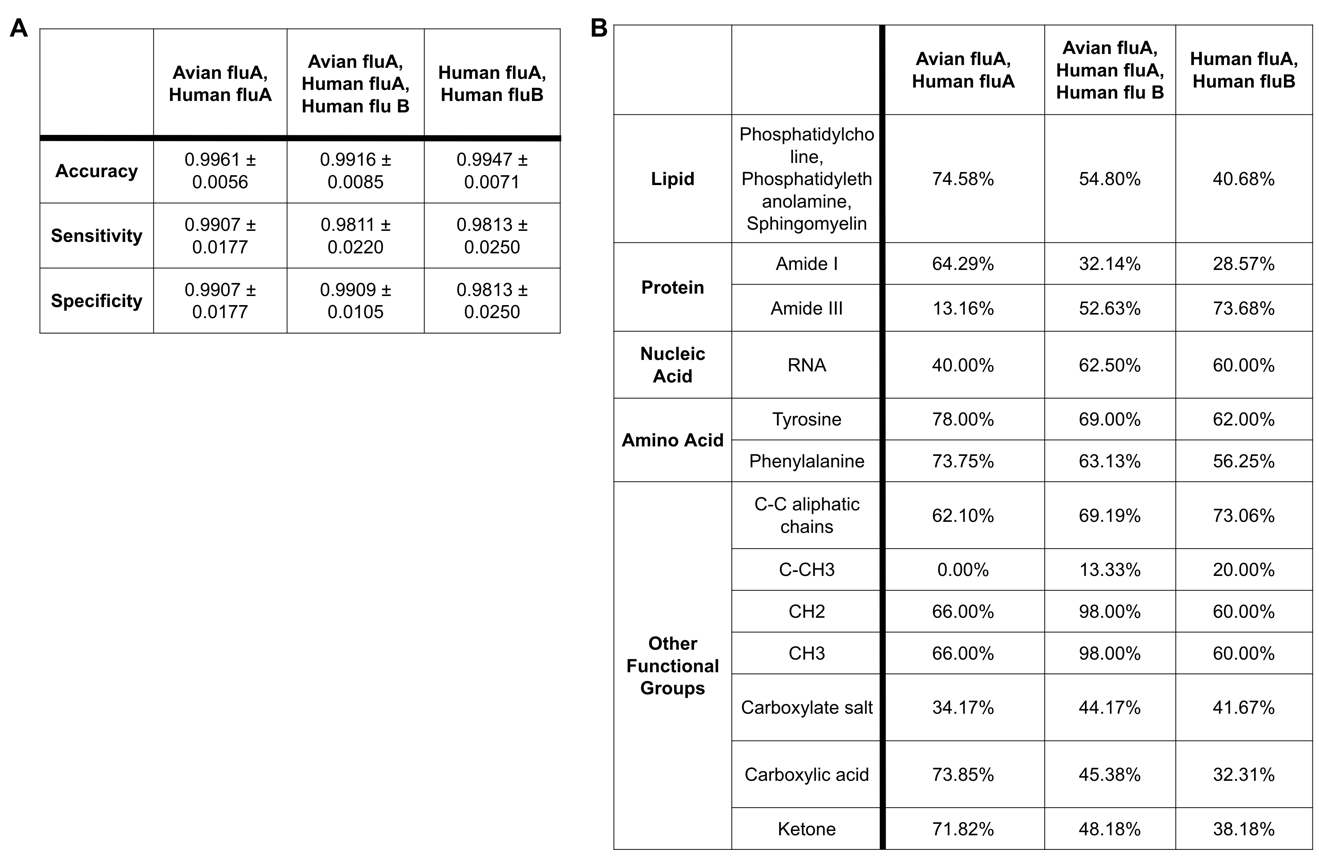

**Fig. S5. A.** The CNN performance on three classification tasks involving avian and human flu viruses (1. Avian Flu A vs. Human Flu A; 2. Avian Flu A, Human Flu A, Human flu B; 3. Human Flu A vs. Human Flu B); **B.** Matching scores between Raman ranges important for each of the three classification tasks using ML and Raman peak ranges of biomolecules.

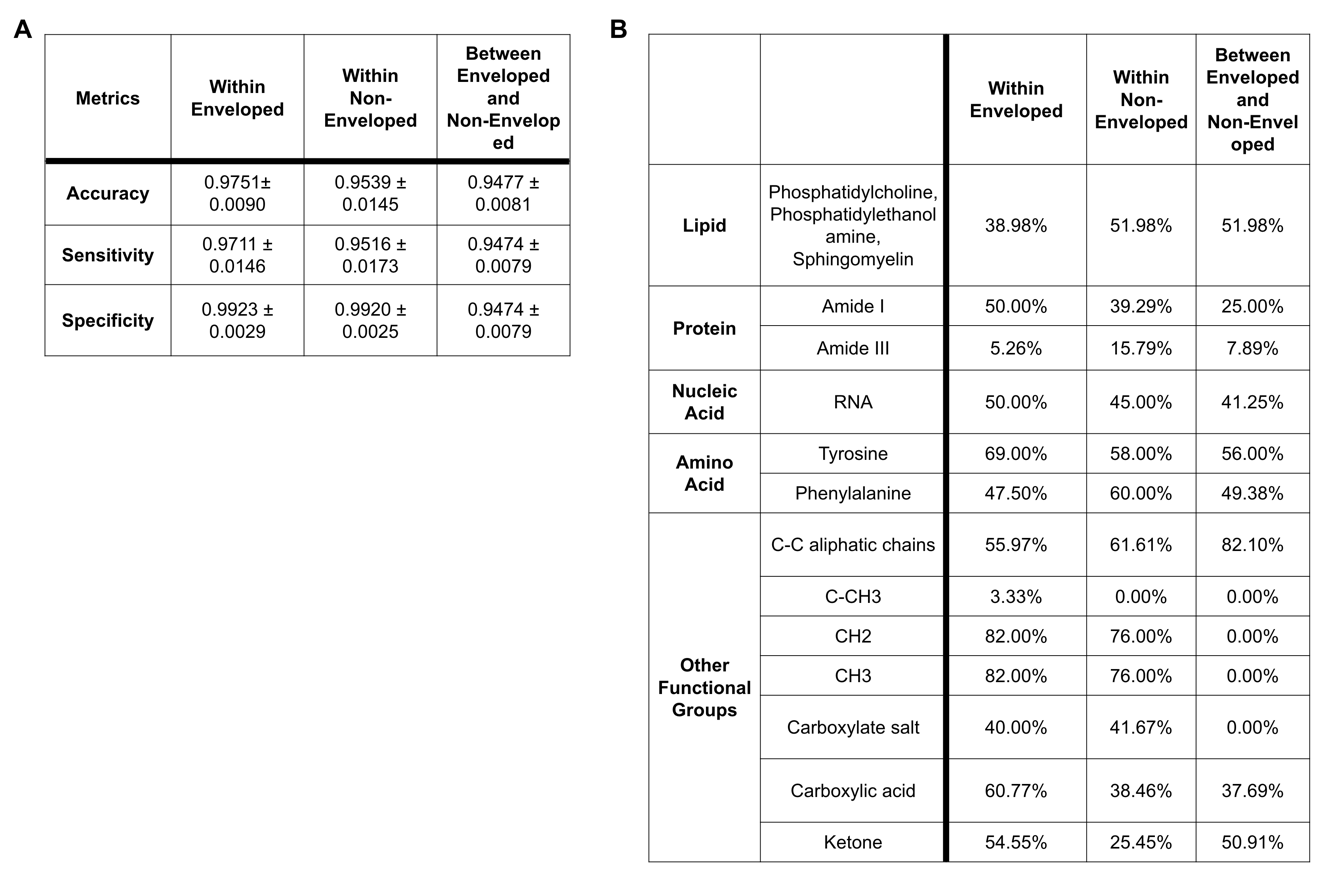

**Fig. S6. A.** The CNN performance on three classification tasks involving enveloped and non-enveloped viruses (1. Classification within enveloped viruses, including Flu A, Flu B, IBV, RSV; 2. Classification within non-enveloped viruses, including Reovirus, Enterovirus, Rhino; 3. Binary classification to identify a virus as either enveloped or non-enveloped; **B.** Matching scores between Raman ranges important for each of the three classification tasks using ML and Raman peak ranges of biomolecules.

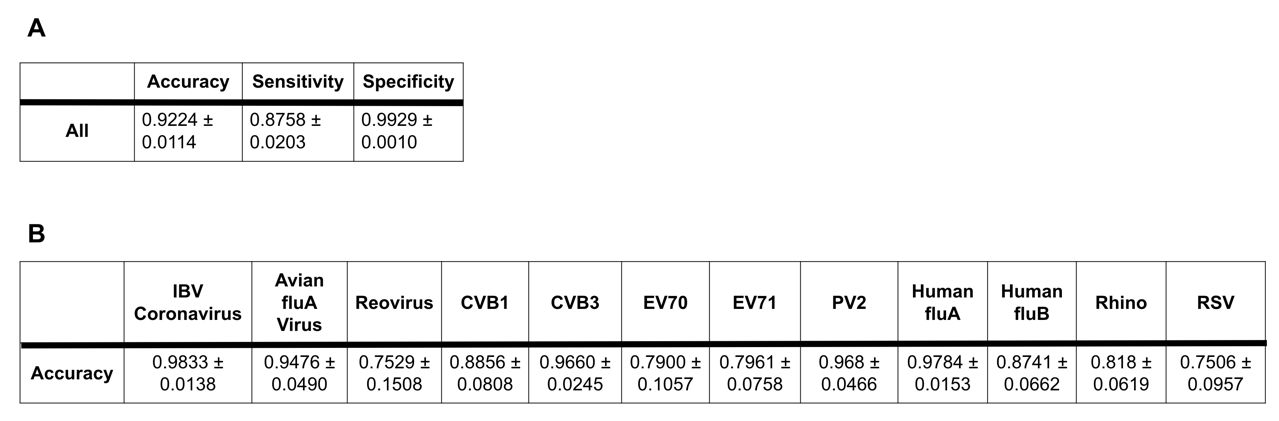

**Fig. S7. A.** The overall CNN performance of classifying / identifying virus type (subtype) among all viruses in our dataset in one classification task; **B.** The classification accuracy for each type of virus, including Avian, Enterovirus and Human Respiratory viruses.

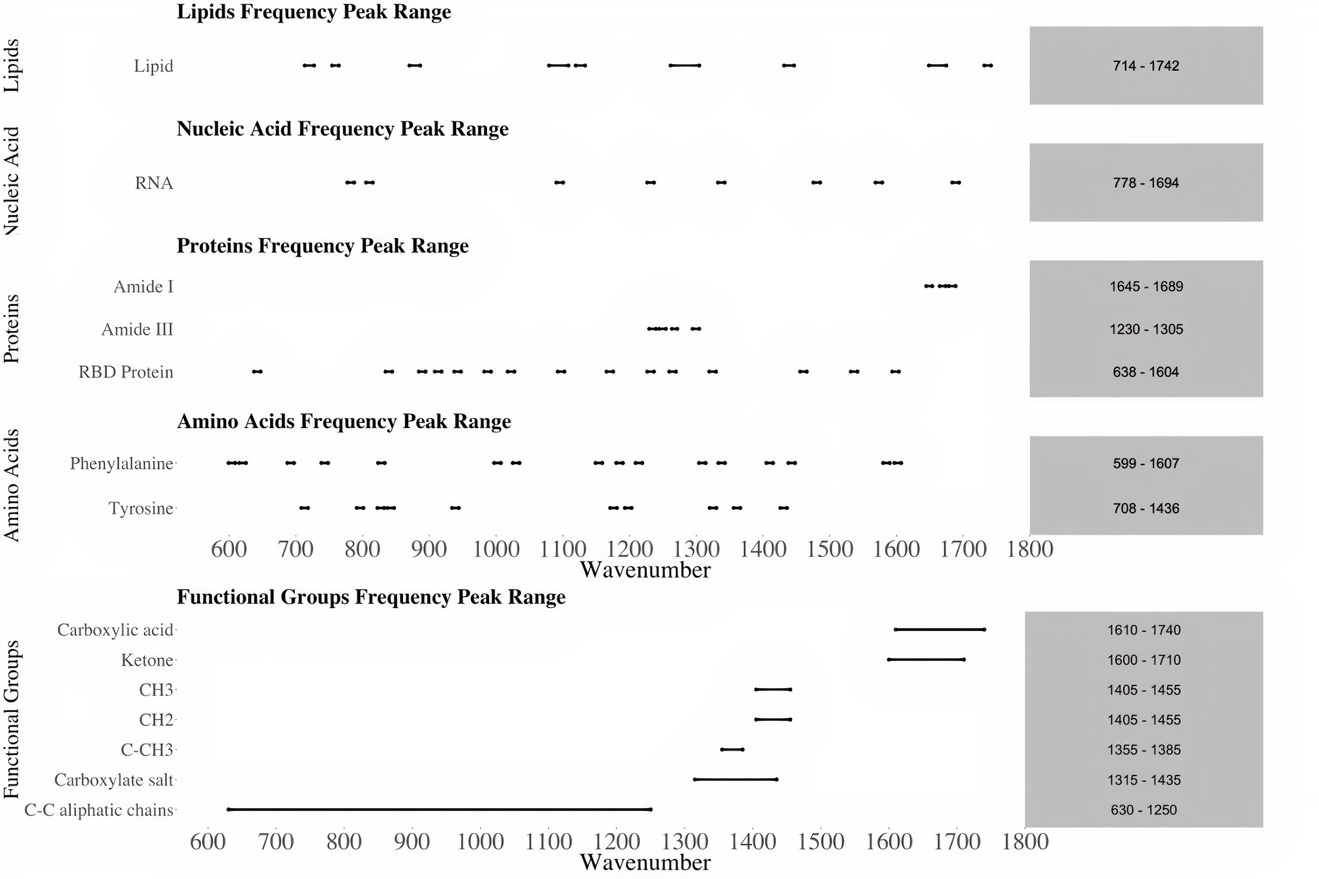

**Fig. S8.** Raman peak ranges of lipids (phosphatidylcholine, phosphatidylethanolamine and sphingomyelin), nucleic acids, proteins, amino acids and other chemical functional groups such as Carboxylic acid and Ketone. These peak ranges are used for matching score calculation in this paper to help us understand what biomolecules or chemical functional groups are important for virus identification tasks using ML.

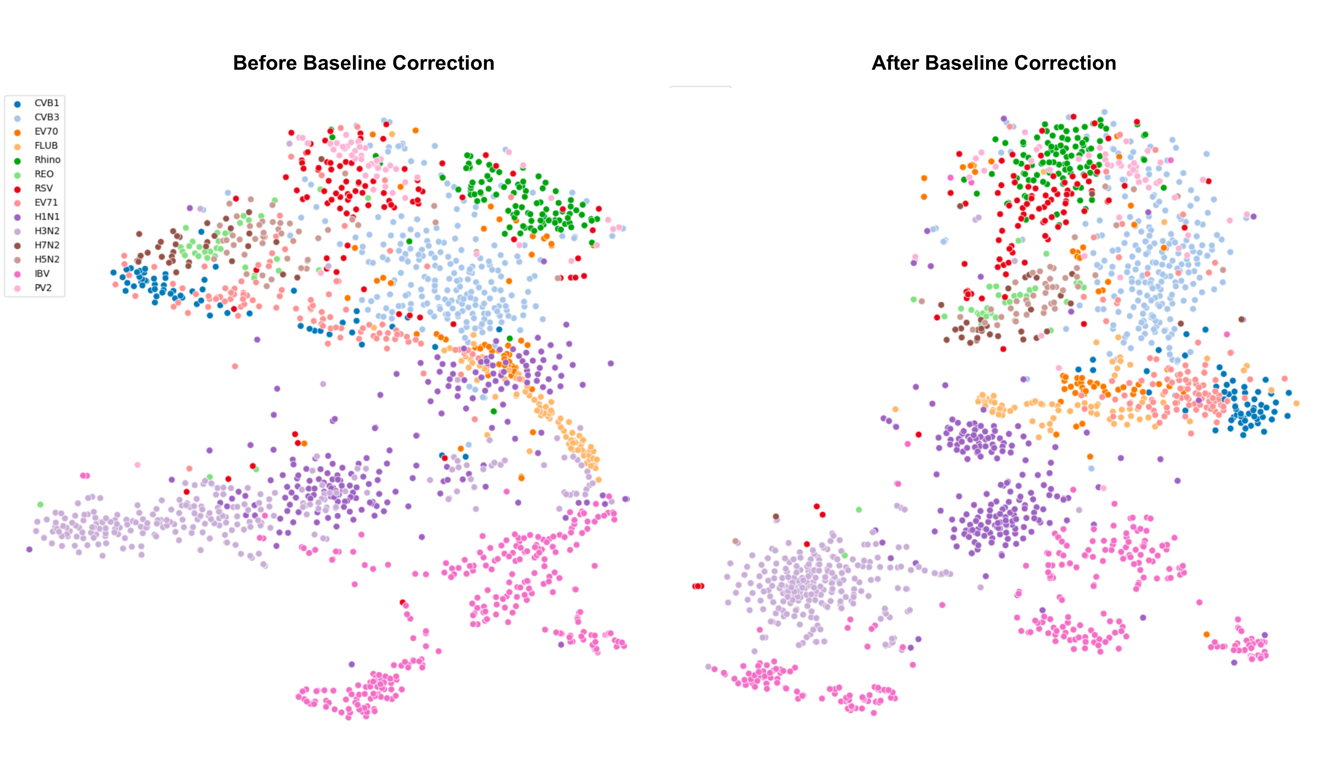

**Fig. S9.** The t-SNE plots of all viruses (Avian virus, Enterovirus and Human Respiratory viruses), before and after baseline correction. Each Raman spectrum is represented by a point in the plots. Observed from the comparison between the two plots, applying baseline correction makes the spectra of virus types (or subtypes) such as H3N2, H7N2, CVB1, RSV, EV71 more distinguishable by pulling tighter each cluster corresponding to spectra of the same virus while pushing the clusters of different viruses further apart.
